# A multiregional assessment of transnational pathways of introduction

**DOI:** 10.1101/2020.11.09.373977

**Authors:** Chris M. McGrannachan, Shyama Pagad, Melodie A. McGeoch

**Affiliations:** Manaaki Whenua – Landcare Research, 231 Morrin Rd, St. Johns, Auckland, 1072, New Zealand; Invasive Species Specialist Group, Species Survival Commission, International Union for Conservation of Nature, University of Auckland, 1072 Auckland, New Zealand; Department of Ecology, Environment and Evolution, School of Life Sciences, La Trobe University, Melbourne 3086, Australia

**Author notes:** Corresponding author: *Chris M McGrannachan.

**Keywords:** Aichi Target 9, alien taxa, Convention on Biological Diversity, introduction event, introduction pathway, propagule pressure

## Abstract

Information on the pathways by which alien taxa are introduced to new regions is vital for prioritising policy and management responses to invasions. However, available datasets are often compiled using disparate methods, making comparison and collation of pathway data difficult. Using a standardised framework for recording and categorising pathway data can help to rectify this problem and provide the information necessary to develop indicators for reporting on alien introductions. We combine the Convention on Biological Diversity’s Pathways Categorisation Scheme (CPC) with data compiled by the Invasive Species Specialist Group (ISSG) to report on multiregional trends on alien introduction pathways over the past 200+ years. We found a significant increase in the documented number of multiregional alien introduction events across all pathways of the CPC’s three hierarchical levels. The ‘escape’ pathway is the most common documented pathway used by alien taxa. Transport stowaways via shipping-related vectors are a rapidly increasing contribution to alien introductions. Most alien introduction events were of unknown pathway origin, highlighting the challenge of information gaps in pathway data and reiterating the need for standardised information-gathering practices. Combining the CPC framework with alien introduction pathways data will standardise pathway information and facilitate the development of global indicators of trends in alien introductions and the pathways they use. These indicators have the potential to inform policy and management strategies for preventing future biological invasions and can be downscaled to national and regional levels that are applicable across taxa and ecosystems.

## Introduction

Expansion and increased intensity of global trade and human movement has exacerbated global species invasions (Essl et al. 2015, Early et al. 2016). Worldwide increases in the number of alien species are likely to continue (Seebens et al. 2017), meaning it is crucial that the pathways by which alien species are transported and introduced to new locations, and how these change in relative importance over time, are identified, understood and better managed (Essl et al. 2015, Chapman et al. 2017). Pathways of introduction are the means by which alien species are transported intentionally or unintentionally outside of their natural geographic range (Richardson et al. 2010, Turbelin et al. 2017). A pathway approach to risk assessment for invasive alien species focuses primarily on identifying introduction pathways to (i) develop early detection and preventative strategies, with the aim to reduce or eliminate the propagule pressure of alien species (Faulkner et al. 2016, Padayachee et al. 2017, Pergl et al. 2017), and (ii) to prioritise investment in managing pathways responsible for the highest propagule loads or particular high risk species (McGeoch et al. 2016). This approach is particularly important in the absence of species-specific data, or when suitable control efforts for individual species are unachievable (Hulme et al. 2008, Padayachee et al. 2017). Accounting for introduction pathways is therefore fundamental for developing relevant management and policy strategies that minimise the introduction, spread and impact of alien species (Hulme et al. 2008).

Efforts to categorize alien species via their pathways of introduction have culminated in the development of a standardised pathway categorisation framework (Harrower et al. 2017). Using this framework, pathways of introduction and spread are classified as intentional or unintentional and encompass three introduction mechanisms: the importation of a commodity, the arrival via a transport vector (through a dispersal corridor resulting from human activity), and the natural spread from a neighbouring region where the species is alien (UNEP 2014, Essl et al. 2015). The foundation of this framework is the six pathway introduction categories (release, escape, transport-contaminant, transport-stowaway, corridor and unaided) originally proposed by Hulme et al. (2008), which encompass 32 specific vectors of introduction (for example, agriculture, horticulture and ship ballast water). This ‘Convention on Biological Diversity (CBD) Pathways Categorisation’ (CPC) (*sensu* Harrower et al. 2017) incorporates standardised terminology and guidelines for pathway categorization and is applicable at a global scale and across different taxonomic groups (Harrower et al. 2017, Tsiamis et al. 2017). The CPC has now been validated by application to alien introductions at national (South Africa; Faulkner et al. 2016), continental (Europe; Pergl et al. 2017, Tsiamis et al. 2017) and global scales (167 cities worldwide; Padayachee et al. 2017). Importantly, the intention of this scheme is, *inter alia*, to assist global reporting as well as country Parties to the CBD to respond to the Strategic Plan for Biodiversity 2011-2020 (UNEP 2014). In particular, this is relevant to achieve and report on Aichi Target 9 by 2020, such that *invasive alien species and pathways are identified and prioritized, priority species are controlled or eradicated and measures are in place to manage pathways to prevent their introduction and establishment* (Convention on Biological Diversity 2010). Whereas monitoring pathways of invasion was not included in the previous global indicator framework for invasive alien species (McGeoch et al. 2010), doing so has now become central to reporting on policy targets for biological invasion (McGeoch and Jetz 2019).

Developing information on pathways introductions using a standardised framework is currently a priority for several reasons. First, preventing the introduction and spread of alien and potentially invasive species is the first line of defence in the management of biological invasions. Managing the early stages of the invasion process (i.e. transport and introduction) that focus on prevention is more cost-effective than reactive, post-introduction management of species (Leung et al. 2002, Rout et al. 2011, Kumschick and Richardson 2013). Nonetheless, management, policy and research that targets the transport and introduction stages of invasion remain relatively underrepresented compared to the invasion stages of establishment and spread (Puth and Post 2005, Early et al. 2016, Chapman et al. 2017).

Second, information on the pathways of species introductions has not, to date, been consolidated into a readily available or accessible form (Saul et al. 2017). Harmonising and identifying discrepancies between data sources is crucial for informing alien species policy and management (Seebens et al. 2020). For example, a recent comparison of European pathway data between the European Alien Species Information Network (EASIN) and the CPC revealed that the pathway subcategories of ~ 5,500 alien species registered with EASIN did not directly align with CPC subcategories (Tsiamis et al. 2017). These types of discrepancies can compound the already high level of uncertainty when identifying and assigning pathways to individual species introductions, particularly for unintentional pathways (e.g. transport-contaminant; transport-stowaway) that may be inadequately documented (Essl et al. 2015).

Third, information on introduction pathways contributes directly to biosecurity policy and regulations, including regulating the criteria for the import and trade of alien species (Burgiel et al. 2006, Leung et al. 2014, Hulme 2015). For example, a blacklist (banned from importation) or whitelist (permitted importation) approach has been adopted by many countries as a response to the global trade in ornamental nursery stock, which is the primary means of introduction of alien plants (Dehnen-Schmutz 2011, Essl et al. 2011, Hulme et al. 2017). Pathway information informs prioritisation of biosecurity interventions by identifying pathways that pose relatively high invasion risk in terms of both propagule load (Brockerhoff et al. 2014) and high risk species (Pergl et al. 2017, Roy et al. 2014) and further informing the development of preventative management strategies and policy at multiple scales (Pyšek et al. 2011, Faulkner et al. 2016). However, few comprehensive pathway-focused policies have been implemented at any administration level, and those that are in place tend to target the release and escape pathways (Essl et al. 2015).

Finally, information on pathway changes over time can, with appropriate modelling and interpretation (McGeoch and Jetz 2019), be used to develop indicators for reporting on alien introduction trends (Rabitsch et al. 2016, Wilson et al. 2018). While the importance of some pathways can remain constant over several decades (e.g. shipping), other pathways (e.g. horticulture) may increase in importance (Ojaveer et al. 2017, Zieritz et al. 2017). These changes may reflect updated legislation for the importation of species, or the increasing global trade of certain commodities (Zieritz et al. 2017, Seebens et al. 2018), and are important for monitoring the effectiveness of biosecurity policy and implementation, such as Aichi Biodiversity Target 9 as well as Sustainable Development Goal 15.8 (Rabitsch et al. 2016).

To date, pathway analysis has been conducted for specific regions (e.g. South Africa, Europe; Faulkner et al. 2016, Pergl et al. 2017), environments (e.g. urban; Padayachee et al. 2017), taxonomic groups (e.g. invertebrates, plant pests; Chapman et al. 2016, Houghton et al. 2016) or specific pathway(s) (Kumschick et al. 2016, Tingley et al. 2018). Although several assessments have shown changes in pathways of invasion over time (Rabitsch et al. 2013, Ojaveer et al. 2017, Zieritz et al. 2017), these are restricted to specific taxonomic groups or geographic locations (but see Rabitsch et al. 2016). Building on these regional and taxon-specific efforts, here we conduct a cross-taxonomic, multiregional analysis of information available on transnational introduction pathways, that incorporates all major groups, environments and pathways, to quantify decadal trends in invasion reported via these pathways since 1800. We use a hierarchical, standard categorisation of pathways (Harrower et al. 2017) so that the results may in future be appropriately modelled, compared, downscaled to regions and countries, and form a baseline for future reporting of trends in invasion pathways. We specifically ask (1) are recorded invasive alien species introductions largely intentional or unintentional? (2) What pathways of introduction and spread are responsible for alien species introductions? (3) What vectors are alien species using to move about?

## Methods

### Data used

Introduction records compiled from the Global Register of Introduced and Invasive Species (GRIIS) by the ISSG were used as the underlying data for the analysis of pathway trends. The GRIIS dataset provides verified and annotated country checklists of alien and invasive species (Pagad et al. 2018). In addition to species names, each record includes taxonomy, the environment/system in which the species occurs, the provenance/origin of the species, evidence of impact (yes/no), date of introduction or first record, type of introduction, pathways of introduction, mechanism of impact, and references for source information. GRIIS Version 2016.2 includes draft checklists for all 196 countries that are party to the CBD.

Data for 18746 introduction events, involving 4832 alien species in 101 countries, and occurring between the years 1300 and 2017, were available and adequate to conduct a pathways assessment (Fig. 1 – map). Here we define an introduction event as a recorded introduction of an alien species in a country outside of its native range. Each introduction event included the date of first introduction or first record of a species and contained data on either all or some of the following information types: (1) introduction being intentional or unintentional (i.e. ‘pathway type’); (2) ‘pathway category’ (escape, release, transport as contaminant or stowaway, corridors, unaided or unknown); (3) vectors (further details of specific vectors within each pathway category, (i.e. ‘pathway subcategory’). The data include Animalia, Bacteria, Chromista, Fungi, Plantae, Protozoa and Virus taxa. The 101 countries cover six regions: Africa, Asia, Europe, North America, Oceania and South America (Figure 2; Table S1). These countries encompass a range of different sizes, development status and climatic regions and thus are geographically representative of global data.

**Figure 1.**
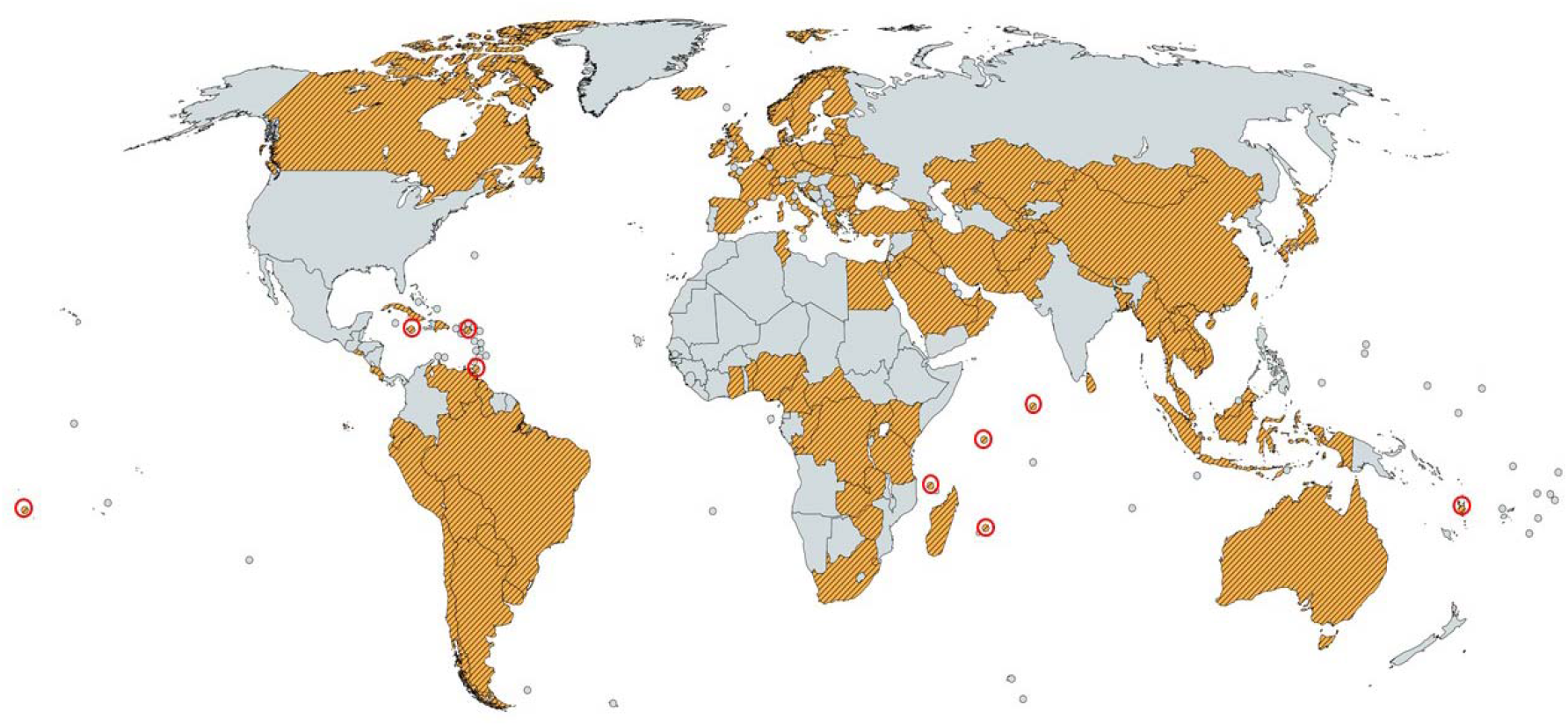
The 101 countries (orange) used to conduct the global pathways assessment. Red open circles indicate small island nations (n = 9) (https://mapchart.net/; accessed 30 July 2019).

**Figure 2.**
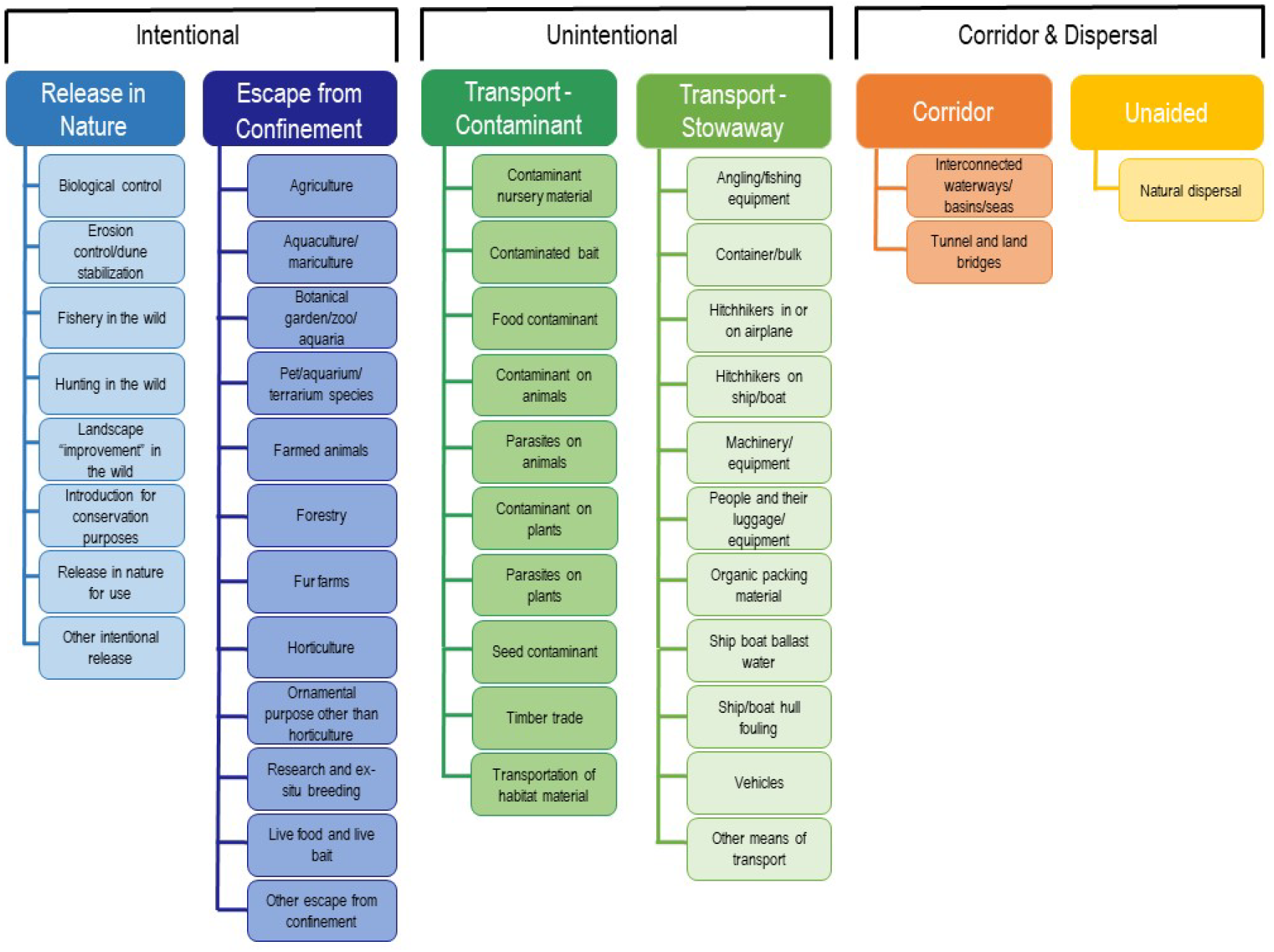
Overview of the hierarchical, standard categorisation of pathways. Six pathway categories and 44 pathway subcategories are broadly categorised into a) intentional transport and introduction of taxa, b) pathways of unintentional introduction and c) pathways by which taxa move to new regions, without direct transportation by humans (i.e. Pathway types). Adapted from Harrower et al. (2017).

A further 5113 species are known to be introduced to the selected 101 countries via known pathways but were not included in analysis as they do not have authoritative information on dates of introduction or first record. These species were therefore excluded and we concentrated on the 4832 species for which the date of introduction in these 101 countries is known. The total number of introduction events currently estimated is ~ 98422, involving ~ 10800 species (including the 5113 species mentioned above). These events, besides known invasive species, include weeds, agricultural pests and diseases, and other non-invasive aliens for which no pathway information or dates of introduction are known.

Information and data on pathways of introduction were extracted during 2016/2017 from sources used to compile national checklists (see Pagad et al. 2018 for information on the general data collation and entry process). Information sources ranged from scientific peer-reviewed literature, databases, reports both published and unpublished and research data. Textual information describing pathways of introduction were documented and then reviewed for categorisation. These categories were inserted into the data collection templates. Because the CPC is relatively new, some of the information from the data sources used pathway terminology that did not fully align with the CBD framework. In these cases, it was necessary to interpret the pathways within the CBD framework, using literature-based pathway information as a guide. This enabled all data to be compiled using the standard categorisation of pathways endorsed by the Parties to the CBD (UNEP 2014). These categories were inserted into the data collection templates.

Information on dates of introduction or first record and information related to the three levels of the pathway hierarchy for the actual introduction event were recorded - pathway type, pathway category and pathway subcategory. Each introduction event was temporally classified using centuries and decades as classifiers (Appendix S1). First introduction records were aggregated by decade beyond 1800. Decadal scales are appropriate because there is often a lag between detection and reporting events. All records prior to 1800 were aggregated as ‘Pre-1800’. Records from the most recent decade were classified as ‘2011 plus’.

### Pathway categorisation

We used the definitions and descriptions of introduction pathways contained in Harrower et al. (2017). This document is the most up-to-date guideline for interpreting the definitions of the CPC and provides examples of the CBD Pathways Categorisation’s application to species information (Harrower et al. 2017). The definitions and descriptions were revised and modified by a panel of experts, using comparisons of the CPC pathway descriptions to descriptions used in (1) the Global Invasive Species Database (GISD), (2) the Delivering Alien Invasive Species Inventories for Europe (DAISIE) database, (3) the Great Britain’s Non-Native Species Information Portal (GBNNSIP) database, and (4) the EASIN information platform (Harrower et al. 2017). Of particular benefit is the distinction between pathway subcategories that appear to overlap. For example, the ‘Contaminant on Plants’ subcategory seemingly overlaps with the ‘Contaminant nursery material’ and ‘Transportation of habitat material’ subcategories. The Harrower et al. (2017) guideline defines and describes the difference between these pathways and treats them in a prescribed order of precedence for category allocation. For example, the ‘Contaminant on plants’ subcategory is defined to contain all contaminants on plants that are not related to the nursery trade, where ‘Contaminated nursery material’ is given precedence over ‘Contaminant on plants’ (Harrower et al. 2017). Despite some shortcomings of the CPC framework, particularly the uncertainty involved in interpreting some subcategories (Faulkner et al. 2020, Pergl et al. 2020), it is a reliable framework with which to report on introduction trends at a transnational level. The CPC framework is still relatively new (2014) and its further development and adoption will facilitate its use as a standardised tool for reporting on alien introductions (Pergl et al. 2020).

### Analysis of trends

For pathway types (i.e. intentional or unintentional introductions), we report trends in terms of both total recorded introduction events for each decade, as well as cumulative introduction events documented between 1800 and 2017. Pathway categories are reported as total number of introduction events per decade for each category. We also report cumulative introduction events for pathway categories, using 1970 as a baseline year. This date was chosen for its comparability with the 1970 baseline used for CBD global biodiversity indicators in Butchart et al. (2010). The dominant pathway subcategories are reported as cumulative introduction events from 1800 to 2017.

We used generalized linear models (negative binomial distribution with log link function) to quantify changes in the recorded number of introduction events over time (introduction events ~ decade). This was conducted at all introduction pathway levels (pathway type, pathway category, pathway subcategory). For subcategories, only the pathways with more than 100 introduction events (n = 18 subcategories) were considered.

## Results

### Pathway types

There was a total of 8172 (43.59%) intentional and 10574 (56.41%) unintentional documented introduction events of alien species across the 101 countries (Table 1). Since 1800, steady and significant increases in both documented intentional and unintentional introduction events have occurred (Table 2; Figure 3a, b). From 1800 to 1900, both pathway types showed similar cumulative increases in introduction events, but with more documented intentional introduction events than unintentional events (Figure 3b, Table 3). Post 1900, the overall number of documented unintentional introductions per decade was higher than intentional introductions (Figure 3a, b; Table 3). Decadal increases in documented introduction events ranged between 5.79 % (1800 – 1810) and 23.15 % (1860 – 1870) for intentional introductions and between 7.19 % (1800 – 1810) and 24.68 % (1890 – 1900) for unintentional introductions (Table 3). The average decadal increase in intentional and unintentional introductions was 13.12 % and 15.29 %, respectively (Table 3). The decade of 1991 – 2000 had more documented introduction events than any other decade in the time series (Figure 3a).

**Figure 3.**
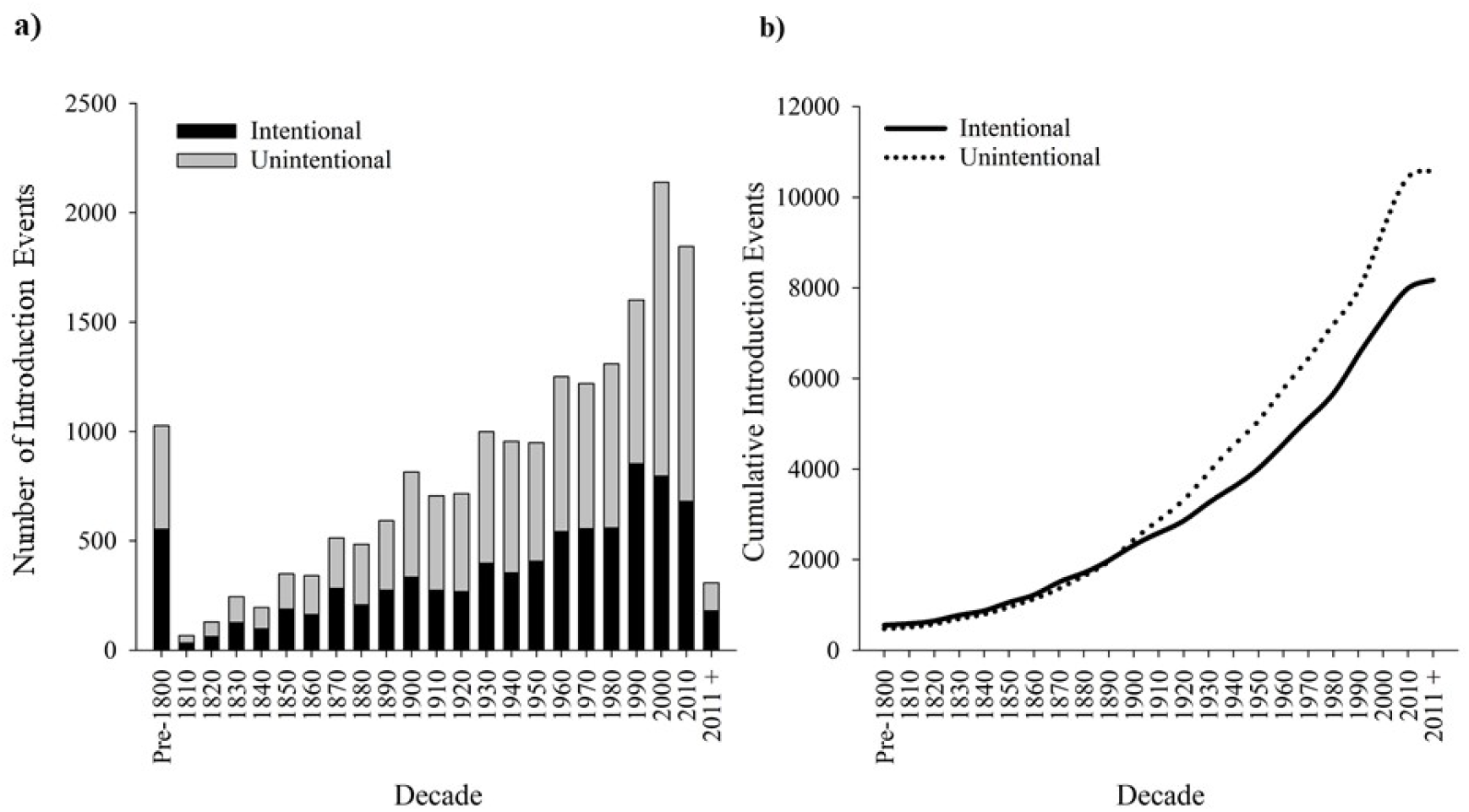
Decadal changes in the documented number of intentional and unintentional introductions of alien species for 101 countries. Trends in introduction events (n = 18746) involving over 4800 alien species are shown as: a) number of documented introduction events, and b) the cumulative number of documented introduction events. An introduction event in this figure represents one species introduced outside of its known native range for the first time and into one of the 101 countries in the pool.

**Table 1:**
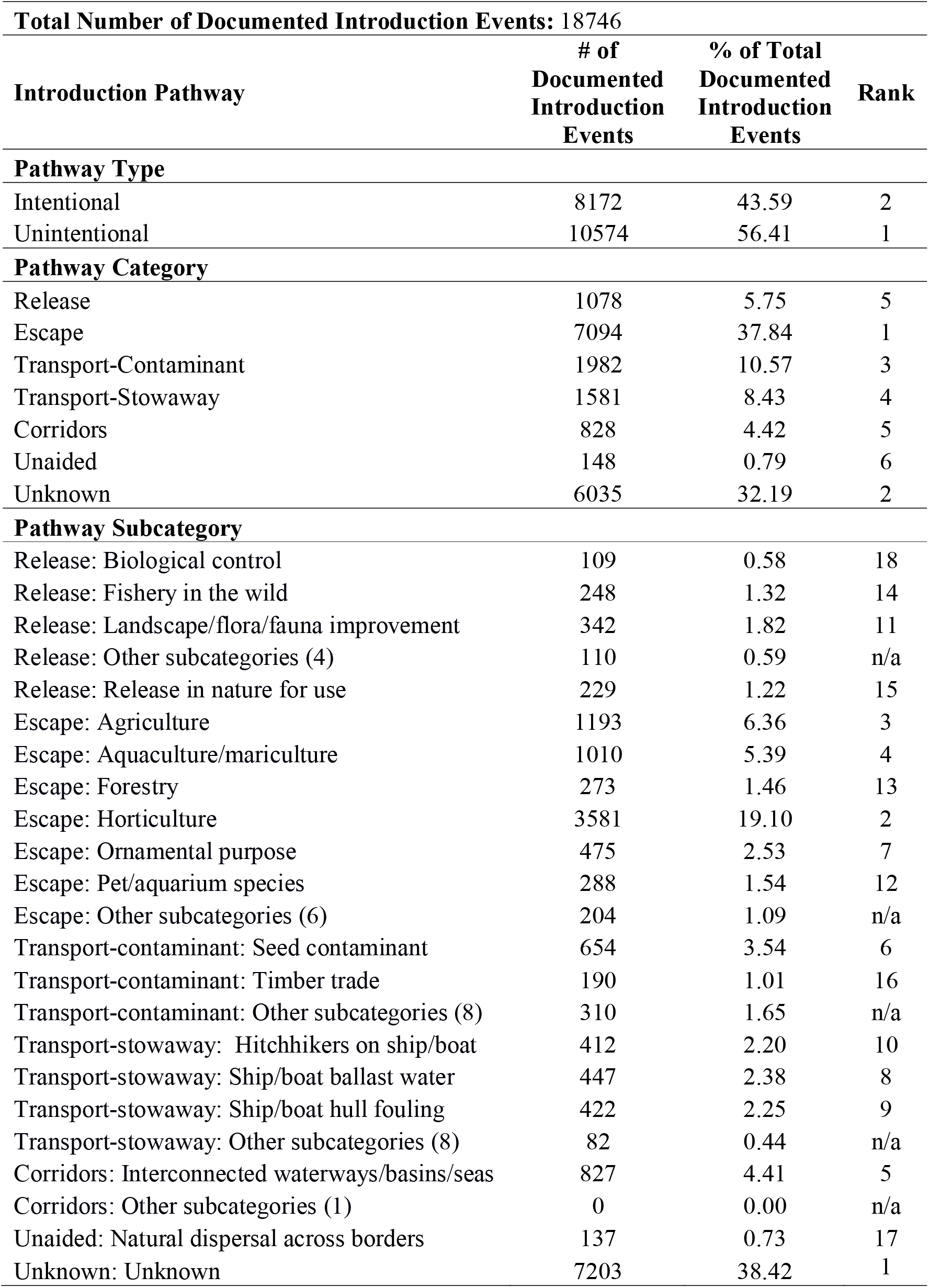
Summary of introduction pathways and their documented introduction events. Bracketed numbers represent the number of subcategories categorised as “other”.

**Table 2.** Decadal increase in documented intentional and unintentional introduction events for the period 1810 to 2011.

| Decade | Documented<br>Intentional<br>Introduction<br>Events | Documented<br>Unintentional<br>Introduction<br>Events | Intentional<br>Decadal<br>Growth<br>(%) | Unintentional<br>Decadal<br>Growth (%) | Difference<br>Between<br>Intentional /<br>Unintentional<br>Decadal<br>Growth<br>Rates |
| --- | --- | --- | --- | --- | --- |
| Pre-1800 | 553 | 473 | n/a | n/a | n/a |
| 1810 | 585 | 507 | 5.79 | 7.19 | 1.40 |
| 1820 | 646 | 575 | 10.43 | 13.41 | 2.98 |
| 1830 | 772 | 693 | 19.50 | 20.52 | 1.02 |
| 1840 | 869 | 790 | 12.56 | 14.00 | 1.43 |
| 1850 | 1056 | 951 | 21.52 | 20.38 | 1.14 |
| 1860 | 1218 | 1129 | 15.34 | 18.72 | 3.38 |
| 1870 | 1500 | 1359 | 23.15 | 20.37 | 2.78 |
| 1880 | 1707 | 1636 | 13.80 | 20.38 | 6.58 |
| 1890 | 1981 | 1953 | 16.05 | 19.38 | 3.32 |
| 1900 | 2314 | 2435 | 16.81 | 24.68 | 7.87 |
| 1910 | 2587 | 2868 | 11.80 | 17.78 | 5.98 |
| 1920 | 2854 | 3316 | 10.32 | 15.62 | 5.30 |
| 1930 | 3251 | 3918 | 13.91 | 18.15 | 4.24 |
| 1940 | 3603 | 4521 | 10.83 | 15.39 | 4.56 |
| 1950 | 4009 | 5064 | 11.27 | 12.01 | 0.74 |
| 1960 | 4550 | 5773 | 13.49 | 14.00 | 0.51 |
| 1970 | 5105 | 6437 | 12.20 | 11.50 | 0.70 |
| 1980 | 5663 | 7188 | 10.93 | 11.67 | 0.74 |
| 1990 | 6516 | 7937 | 15.06 | 10.42 | 4.64 |
| 2000 | 7313 | 9279 | 12.23 | 16.91 | 4.68 |
| 2010 | 7993 | 10445 | 9.30 | 12.57 | 3.27 |
| 2011 > | 8172 | 10574 | 2.24 | 1.24 | 1.00 |
| **Average** | n/a | n/a | **13.12** | **15.29** | **3.10** |
| **Std Dev.** | n/a | n/a | **4.70** | **5.25** | **2.17** |

**Table 3.**
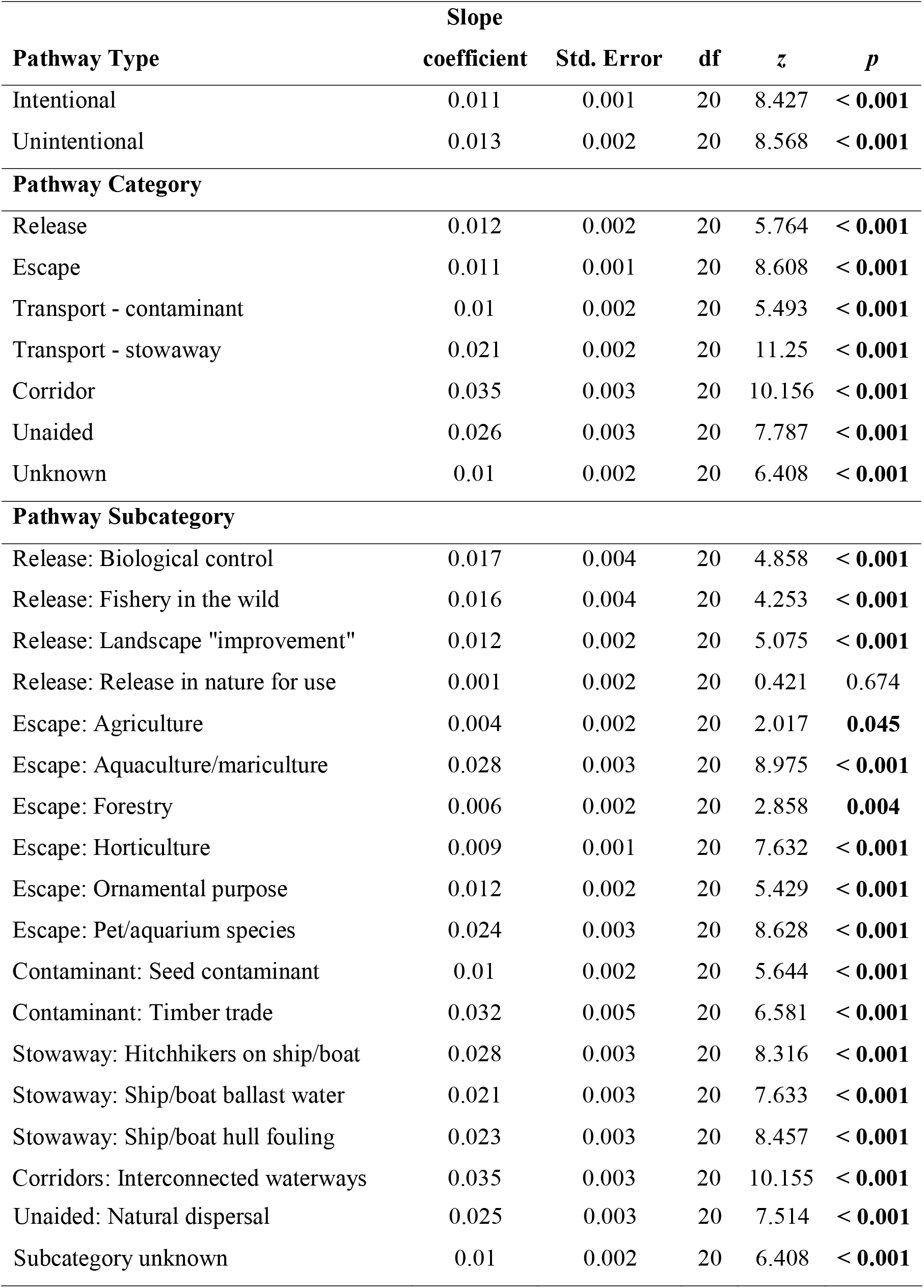
Trends in recoded introduction events by pathway across the period 1800 to 2017. Generalized linear model results (family = negative binomial, link = log). Significant *p* values (*p* < 0.05) shown in bold.

### Pathway categories

The documented number of introduction events for each pathway category has increased significantly per decade since 1800 (Table 2; Figure 4a). The ‘escape’ pathway is the most prevalent pathway by which species introductions are known to occur (37.84 %), followed by ‘unknown’ pathway introductions (32.19 %; Table 1; Figure 4a). Post 1970 trends show both escape and unknown pathways increased dramatically in cumulative number of introduction events, with 3177 and 2350 additional events, respectively, occurring between 1970 and 2017 (Figure 4b). This is equivalent to 81.38 % (escape) and 58.14 % (unknown) of the total number of pre-1970 documented introduction events. The remaining five pathway categories had fewer cumulative introduction events compared to escape and unknown pathways, the highest being ‘transport-contaminant’ (1982 events by 2017) and the lowest ‘unaided’ (148 events by 2017; Figure 4c). Of these five pathways, ‘transport-stowaway’ showed the steepest cumulative increase in introduction events post 1970 (Figure 4c).

**Figure 4.**
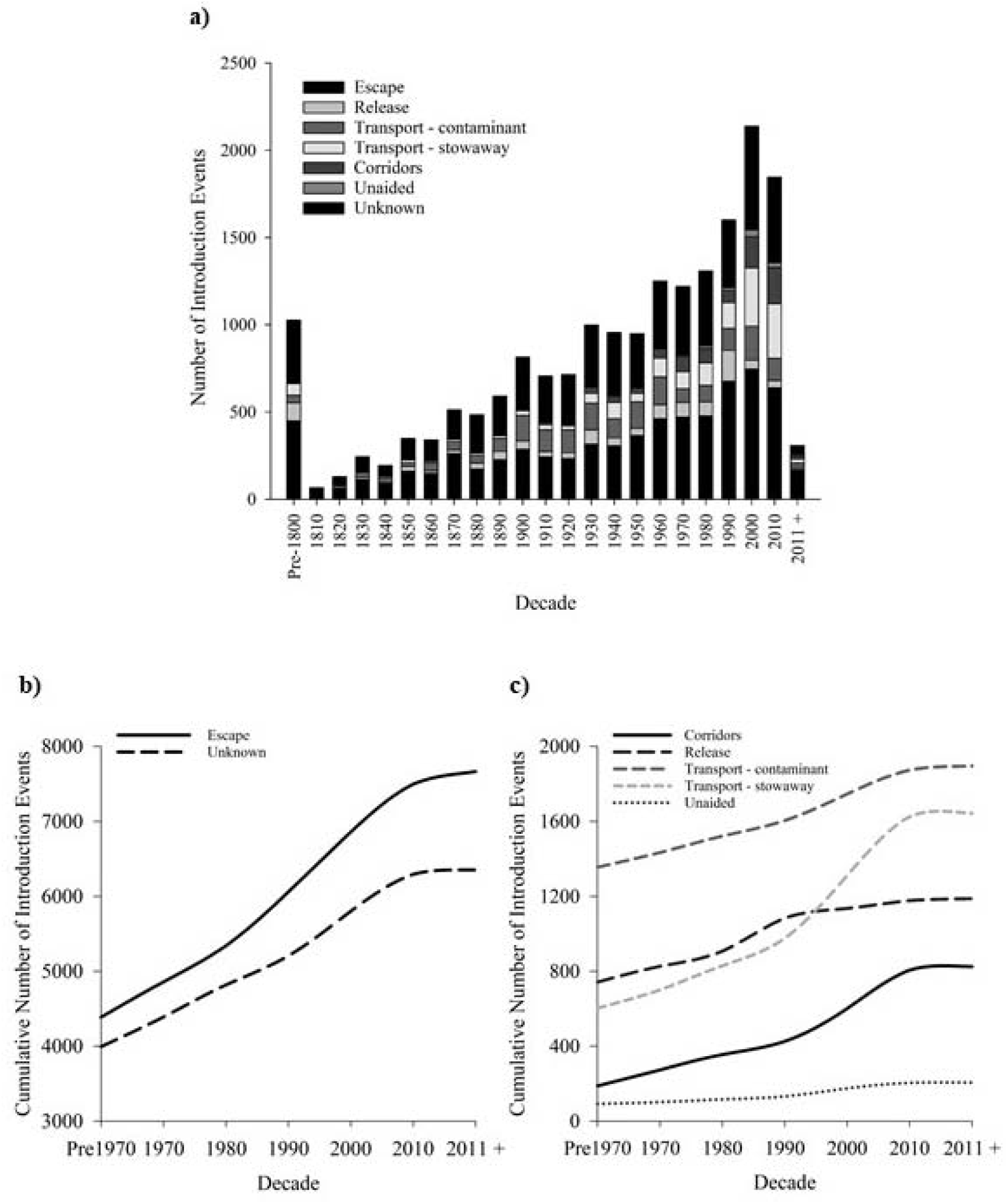
Changes in the six main pathway categories (as well as the number of introductions via unknown pathways). (a) The documented number of introduction events (n = 18746) of alien species per decade since 1800 for 101 countries. (b-c) Cumulative number of documented introduction events by pathway since 1970 (note different scaling on y-axes).

### Pathway subcategories

The 18 pathway subcategories with more than 100 introduction events since 1800 ranged from 109 to 7203 records (Table 1). The top 18 subcategories were representative of all pathway categories. ‘Unknown’ was the pathway subcategory associated with most documented introduction events (7203; 38.42 %), followed by three subcategories from the escape pathway: ‘horticulture’ (3581; 19.10 %), ‘agriculture’ (1193; 6.36 %) and ‘aquaculture/mariculture’ (1010; 5.39 %; Table 1; Figure 5a). Many of the subcategories showed sharp rates of increase, particularly from the beginning of the twentieth century, including ‘hitchhikers on ships’, ‘ship ballast water’, ‘ship hull fouling’ and ‘interconnected waterways’ (Figure 5b, c; Figure S1a). In comparison, most subcategories from the escape pathway (except horticulture) had slower cumulative introduction rates, including ‘agriculture’, ‘aquaculture/mariculture’, ‘forestry’, ‘ornamental purpose other than horticulture’ and ‘pet/aquarium species’. All but one subcategory (‘release in nature for use’) significantly increased in introduction events per decade since 1800 (Table 3).

**Figure 5.**
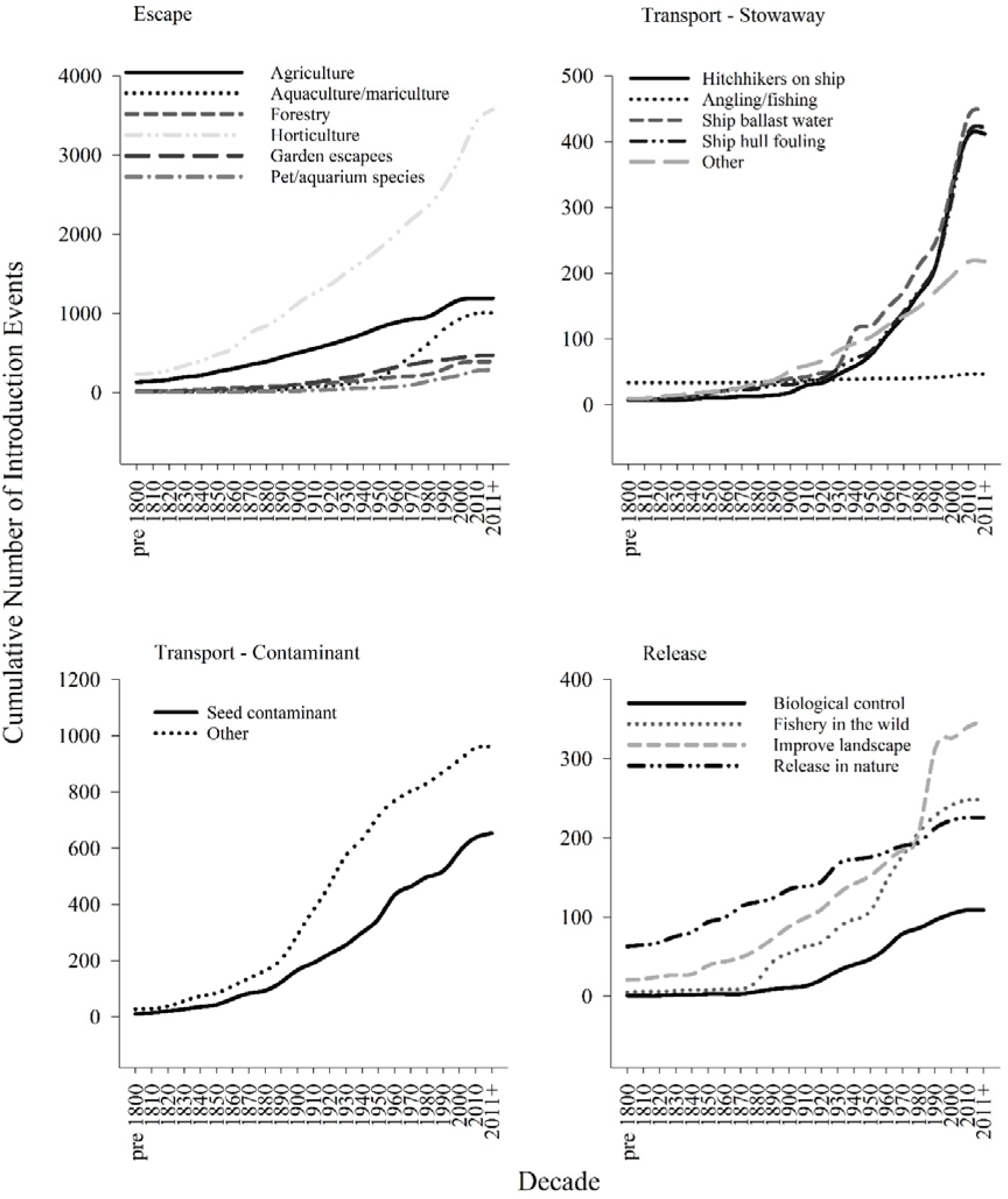
Changes in the dominant pathway subcategories across decades. Cumulative number of documented introduction events (note different scales on y-axes). The 18 pathway subcategories shown are those with most (> 100) introduction events (see Figure S1 for Corridors: interconnected waterways/basins/seas’, ‘unaided: natural dispersal across borders’ and ‘unknown’ pathway subcategories not shown).

## Discussion

We used the CBD pathways categorisation framework and a multiregional dataset encompassing a range of taxonomic groups to report on decadal changes in introduction pathways reported for alien species since 1800. We highlighted the significant increase of documented events for almost every pathway at each of the three hierarchical levels of the CPC. Unintentional introductions have increased over intentional introductions since the beginning of the twentieth century. However, ‘Escape’ – an intentional pathway - is the most common pathway category documented, particularly for vectors related to plant and aquatic cultivation. This shows that intentional pathway vectors are still an important source of alien introductions. The vast majority of documented introduction events, however, have an unknown vector (38.42 %), which emphasises the high level of uncertainty involved in categorising and managing alien species introduction pathways.

### Accidental and deliberate introduction events

Prior to the twentieth century, the cumulative rate of increase for both intentional and unintentional introduction events documented were virtually identical (Figure 3b). The beginning of the twentieth century saw unintentional surpass intentional introductions, a trend that has continued up to the present. The increase in unintentional introductions is likely due to the rise in international trade, which is widely acknowledged as an important factor in allowing alien species to successfully establish in novel geographic regions (Levine and D’Antonio 2003, Perrings et al. 2005, Yemshanov et al. 2012, Chapman et al. 2017). In particular, the accidental transport of inconspicuous taxa, such as fungi, microorganisms, pathogens and invertebrates are often associated with global trade, including live plant imports and importation via shipping (Brockerhoff and Liebhold 2017, Chapman et al. 2017, Okabe et al. 2017). Studies from multiple regions including Europe (Chapman et al. 2017, Pergl et al. 2017, Saul et al. 2017), Asia (Hong et al. 2012, Okabe et al. 2017), the US (Liebhold et al. 2012) and the Antarctic (Osyczka et al. 2012, Houghton et al. 2016) have found that these taxonomic groups are more often associated with unintentional pathways. Increases in trade volume and the subsequent rise in accidental introductions of alien species may counteract existing national biosecurity and phytosanitary measures (Brockerhoff and Liebhold 2017). It is therefore important to improve measures for monitoring unintentional introduction pathways to effectively address the ongoing occurrence of accidental alien introductions. Interestingly, although unintentional introductions surpassed intentional introductions, escape (an intentional introduction pathway category) had most associated introduction events (excluding unknown events). This highlights that the prevention and management of intentional introductions are of equal importance to those of unintentional introductions, especially given that the impact realised by alien taxa has been associated more frequently with intentional than unintentional introductions (Pergl et al. 2017).

### Introduction pathway categories and their vectors

Our findings corroborate previous studies of alien introduction pathways in several ways. First, ‘escape’ is overall the most common documented pathway category by which alien species are introduced (Turbelin et al. 2017). Second, ‘transport-stowaway’ is becoming an increasingly important introduction pathway, particularly for marine stowaways (Zieritz et al. 2017). Finally, records of introduction events via unknown pathways are prevalent in existing databases and presents an ongoing problem for assessing alien introductions (Katsanevakis and Moustakas 2018). Our global perspective takes into consideration alien species from multiple taxonomic groups but supports similar findings from studies focussing on specific taxonomic groups or regions.

Escape was the most prevalent pathway, with records almost doubling between 1970 and the present (Figure 4b). Escape has been identified as the most frequent introduction pathway across all taxa at global (Turbelin et al. 2017) and national (South Africa; Faulkner et al. 2016) scales, for plants at country- (Czech republic; Pyšek et al. 2011; USA; Lehan et al. 2013) and city-scales (Padayachee et al. 2017) and for both plants (Pergl et al. 2017) and vertebrates in Europe (Saul et al. 2017, Roy et al. 2019) and globally (Saul et al. 2017, van Kleunen et al. 2018). Our results corroborate these findings at a multiregional level and emphasise the ongoing need for better containment procedures and greater public awareness of the risks involving escaped organisms, particularly ornamental plants (Ricciardi et al. 2017, Saul et al. 2017, van Kleunen et al. 2018).

Horticulture is the most important vector of alien plant introductions (Turbelin et al. 2017, van Kleunen et al. 2018) and was the pathway subcategory with the largest and fastest increase in introduction events (Figure 5a). Agriculture was the second most important pathway subcategory and is also recognised as an important contributor to alien plant introductions (Mack and Erneberg 2002, Richardson et al. 2003). Both horticulture and agriculture are pathway subcategories specific to plants (Harrower et al. 2017) and their combined, high proportion of recorded introductions in the dataset (see Table 1) supports previous studies that show escape (from horticulture or agriculture) is an important pathway for plants.

The importance of escape as an introduction pathway for faunal species is reflected by the high number of introduction events attributed to escape from aquaculture/mariculture (e.g. fish farms) compared with the pet/aquarium trade. Aquaculture/mariculture had the third most introduction events, while records attributed to the pet/aquarium trade remained relatively stable across the assessed time-period (Figure 5a). Aquaculture was found to be the highest contributing pathway to freshwater alien species introductions in Europe (Nunes et al. 2015) and an important pathway for alien invasions of European seas (Nunes et al. 2014). The ecological impacts of invasion via aquaculture can be severe (Naylor et al. 2001, Keller et al. 2011) and given the aquaculture sector is one of the fastest growing global primary industries (Teletchea and Fontaine 2014), it is also likely that alien introductions via this pathway will continue to rise.

Vectors of the transport-stowaway category were among those with largest growth in alien introductions since 1970 (Figure 5b). In particular, there was a sharp rise in the post-1970 introduction of marine stowaways as hitchhikers on ships, in ship ballast water or as ship hull fouling, which saw 67 %, 62 % and 67 %, respectively. The importance of marine/aquatic pathways is also reflected in the sharp rise in introduction events by interconnected waterways since 1970 (Figure S1a). These increases in alien introductions are likely due to the continued expansion of tourism and international shipping (Early et al. 2016, Turbelin et al. 2017). The introduction of marine and freshwater alien taxa via shipping-related transport has been confirmed as an important source of ongoing propagule pressure in many parts of the world, including the Mediterranean region, Northwest Europe, the Northeast Pacific and Australia (Tingley et al. 2017, Zieritz et al. 2017, Anil and Krishnamurthy 2018).

A key challenge in attempting to decipher trends in alien introductions is uncertainty in the specific pathways used by species (Katsanevakis et al. 2013, Essl et al. 2015). This is particularly problematic for unintentional introductions via transport contaminants or stowaways, and for smaller organisms such as marine invertebrates that are at a higher risk of going unnoticed or undocumented (Essl et al. 2015, Ojaveer et al. 2017, Zieritz et al. 2017). The results shown here demonstrate the problem clearly: the total number of introduction events where a pathway category was unknown far exceeded all other known pathway categories (Figure 4b-c). The exception to this was the ‘escape’ pathway, an intentional pathway category that surpassed the number of unknown introduction events (Figure 4b). Furthermore, ‘unknown’ was the highest-ranked pathway subcategory in terms of the number of introduction events and was almost double that of the second-ranked subcategory (Horticulture; Table 1). These results corroborate previous studies that have demonstrated and highlighted the risk that uncertainty poses to introduction pathway datasets and trends (Zenetos et al. 2017, Katsanevakis and Moustakas 2018).

There are several reasons why uncertainty in pathway identification and trends occurs. Often, the lack of historical introduction records (i.e. pre-mid twentieth century; Ojaveer et al. 2017) can result in gaps in datasets that can particularly impact the interpretation of introduction temporal trends (McGeoch et al. 2010, Katsanevakis et al. 2013, Galil et al. 2018). Usually this occurs due to decreased scientific effort or reduced awareness of the need to record alien species introductions (Ojaveer et al. 2017).

In many cases, multiple pathways are equally tenable as the cause of alien species introductions to a new region (Minchin 2007). This makes assigning the correct pathway difficult and decisions may be entirely based on the interpretations or assumptions of assessors (Zenetos et al. 2012). In other instances, the species’ ecology may be used to infer an introduction pathway (Zenetos et al. 2012). A representative example of this is the introduction of marine species into the Mediterranean Sea via the Suez Canal. Several pathway vectors could feasibly be responsible for new introductions into the Mediterranean, including species as hitchhikers on ships, through ship ballast water or hull fouling, or through natural, unaided dispersal (Katsanevakis et al. 2013). These types of uncertainty can potentially over- or under-emphasise certain pathways, causing trends to be misrepresented at both global and regional scales. Using a confidence score in allocating pathways may provide a cautionary approach to the compilation of pathway data that helps identify which species, pathways or regions are particularly susceptible to uncertainty (Essl et al. 2015). A focus on improving monitoring of these identified species, pathways or regions may aid efforts to alleviate uncertainty in pathway data. Confidence scores have been successfully integrated into other alien-focused, standardised frameworks, such as the Environmental Impact Classification of Alien Taxa (EICAT; Blackburn et al. 2014, Hawkins et al. 2015) and have recently been used in assessing alien introduction pathways in Europe (Pergl et al. 2020).

The compilation of pathway data from multiple countries or regions can also be a source of uncertainty. Data is often unavailable in many countries, due to a lack of adequate monitoring, data collection efforts or funding (Latombe et al. 2017). Compiling data at national or regional levels usually requires a well-established network of contacts and managing these networks can expend a great deal of time and effort (Zenetos et al. 2017). Furthermore, having multiple pathway data sources will result in multiple ways in which the data is formatted, leading to discrepancies between data. Enacting a standardised framework such as the CPC to filter and arrange pathway data will ensure that trends in introductions of alien species are reported accurately. This is crucial if pathways of introduction are to be considered as an accurate indicator for alien species invasions (Wilson et al. 2018). Given that trends in pathway introductions change over time and across regions, the accuracy and standardisation (or lack thereof) of data can greatly benefit or hinder monitoring and biosecurity efforts (McGeoch et al. 2016, Latombe et al. 2019).

Developing indicators from standardised pathway data is necessary for accurate reporting of alien introduction trends. These indicators can then be used to identify the shortcomings in invasive alien species management and policy targets and help improve legislation for dealing with biological invasions (McGeoch et al. 2010, Hulme 2015). Predictive tools such as risk assessments and horizon scanning can incorporate pathway indicators to better estimate the susceptibility of regions to invasion and identify those species that will pose the greatest introduction threat (Hulme 2015, Rabitsch et al. 2016). The continual input of new pathway data will be needed to ensure that indicators remain up to date and to prevent policy decisions relying on historical pathway patterns (Latombe et al. 2019). Given that the Strategic Plan for Biodiversity 2011-2020 is coming to an end, and the 2021-2030 phase is about to begin, the development and testing of pathway indicators for tracking invasive alien species trends becomes increasingly urgent (Rabitsch et al. 2016).

## Conclusions

We propose that the CBD Pathway Categorisation framework is a suitable tool for providing standardised information on alien introduction pathways. This information can then be used to report on pathway trends and their changes across time, taxa, habitats and geographic scales. However, the high number of cases where introduction pathways are unknown will remain a significant challenge to the reporting and documentation of alien introductions (Latombe et al. 2019). Despite this, the CPC framework can enable countries to improve recording and reporting of alien introductions and assist in developing strategies to reduce the impacts of alien introductions beyond the Strategic Plan for Biodiversity 2011-2020.

## Acknowledgements

MAM acknowledges support from the Australian Research Council (DP200101680).

