## Supplementary material for "A multiregional assessment of transnational pathways of introduction": Figure S1

**Supporting Information**

**Figure S1.** Cumulative introduction events between 1800 and 2017 for a) interconnected waterways and natural dispersal pathway subcategories and b) unknown pathway subcategories.

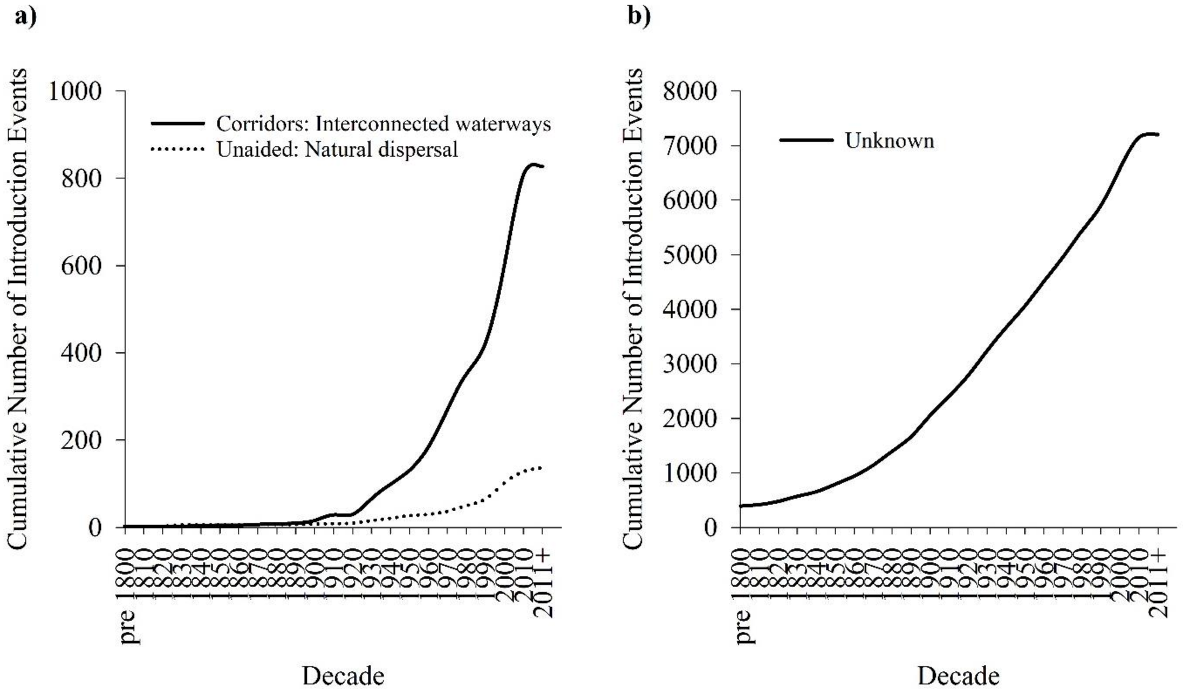
