## Supplementary material for "A multiregional assessment of transnational pathways of introduction": Table S1

**Supporting Information**

**Table S1.** List of 101 countries used in the analysis of introduction pathway trends.

| COUNTRY NAME | COUNTRY NAME | COUNTRY NAME | COUNTRY NAME |
|---|---|---|---|
| Afghanistan | Czech Republic | Kazakhstan | Slovakia |
| Albania | Democratic Republic of Congo | Kenya | South Africa |
| Argentina | Denmark | Lao People's Democratic Republic | Spain |
| Armenia | Dominican Republic | Latvia | Sri Lanka |
| Australia | Ecuador | Lithuania | St. Kitts and Nevis |
| Bangladesh | Egypt | Madagascar | Swaziland |
| Belarus | El Salvador | Malaysia | Sweden |
| Belgium | Estonia | Maldives | Switzerland |
| Benin | Finland | Mauritius | Taiwan |
| Bhutan | France | Mongolia | Tajikistan |
| Bolivia | Georgia | Myanmar | Tanzania |
| Brazil | Germany | Nepal | Thailand |
| Bulgaria | Ghana | Netherlands | Trinidad and Tobago |
| Cambodia | Greece | Nigeria | Tunisia |
| Cameroon | Guyana | Norway | Turkey |
| Canada | Iceland | Oman | Uganda |
| Central African Republic, the Congo | Indonesia | Pakistan | Ukraine |
| Chile | Iraq | Paraguay | United Arab Emirates |
| China | Ireland | Peru | United Kingdom |

| COUNTRY NAME | COUNTRY NAME | COUNTRY NAME | COUNTRY NAME |
|---|---|---|---|
| Comoros | Islamic Republic of Iran | Poland | Uruguay |
| Cook Islands | Israel | Romania | Uzbekistan |
| Costa Rica | Italy | Rwanda | Vanuatu |
| Croatia | Jamaica | Saudi Arabia | Venezuela |
| Cuba | Japan | Seychelles | Viet Nam |
| Cyprus | Jordan | Singapore | Zambia |
| | | | Zimbabwe |
